# Endothelial Cells stimulate proliferation of CD140a sorted human Glial Progenitor Cells and their specification towards astrocytic lineage

**DOI:** 10.1101/2020.03.06.981456

**Authors:** Asmita Dasgupta

## Abstract

Homeostasis in stem cell niche is established by fine-tuning between interacting signaling pathways of all resident cell types that constitute the niche. In this context, the exact contribution of human endothelial cells in biology of PDGFRα positive human glial progenitor cells (hGPCs) in niche environment is not well understood. Towards the same, insert co-culture system with human umbilical vein endothelial cells (HUVECs) has been adopted. In co-culture with HUVECs, under proliferative condition, hGPCs show increased proliferation and sphere formation, while under differentiating condition hGPCs show increased differentiation to astrocytes with concomitant decrease in differentiation to oligodendrocytes with respect to no co-culture controls. Transcript assay for selected humoral factors reveals bone morphogenic proteins (BMPs), endothelin1, growth arrest specific 6 and interleukin 6 to be in higher abundance in HUVECS than hGPCs, of which BMP4 transcripts were most abundant, indicating possibilities of its being the key mediator of endothelial mediated effects. Concurrently, noggin effectively attenuates HUVEC mediated astrocytic differentiation of PDGFRα sorted fetal hGPCs The results have implication towards safety of transplantation therapies with PDGFRα sorted fetal hGPCs. It is postulated that proliferation and differentiation response as seen in this co-culture system could have defining implications towards the genesis of glioma.

## Introduction

Among the cells that constitute the niche, vascular system forms an integral part^1,2^. Vascular support of neuronal recruitment mediated by BDNF and active vascular recruitment and remodeling by the neurons mediated by VEGF is essential for adult neurogenesis^3,4,5^. Vascular endothelial cells have been demonstrated to stimulate self-renewal by activating Notch and Hes1 in neural stem cells (NSCs) and expand neurogenesis *in vitro* murine systems^6^. Relatively less is known about vascular contribution to proliferation, maintenance and migration of the bi-potential oligodendrocyte-astrocyte precursors, the glial progenitor cell (GPC)^7,8^ and even less so particularly for human species. The GPCs are mainly identified by either their expression of platelet-derived growth factor receptor-alpha (PDGFRα)^9,10,11^ or by immune-selection by A2B5 monoclonal antibody^12,13,14^. These cells mature to express transcription factors Sox10, Olig-1 and Olig-2, followed by expression of O4 and NG-2 proteoglycan as well as cyclic nucleotide phosphodiesterase and myelin basic protein to form committed oligodendrocytes^15^.

It has been known that NSC-endothelial contacts in sub-ventricular zone (SVZ) are frequently devoid of astrocytic endfeet or pericyte sheath and thus are unusually permeable, allowing NSCs to be constantly exposed to diffusible endothelial derived factors. *In vitro* murine systems, have shown glial differentiation of cortical progenitors to be regulated by the vascular endothelial cells mediated by bone morphogenic proteins (BMPs)^16^. Endothelial cells are also known to promote survival and proliferation of oligodendrocyte precursor cells by Akt and Src signaling pathways^17^. Recently, Olig2+ PDGFRα+ human OPCs during 18-24gw have been shown to be associated to vasculature which they apparently use as scaffolds for migration^18^. Co-culture of NSCs in direct contact with endothelial cells is known to transiently delay spontaneous apoptosis without modifying their overall proliferation by reversible increase in number of Lex expressing quiescent progenitors mediated by BMP/Smad signaling. This is known to stimulate their differentiation mainly into astrocytes, slightly to neurons and mostly reduce the progenitor differentiation into oligodendrocytes in comparison to adherent cultures on poly-ornithine^19,20^.

Since the vascular support for neuronal recruitment as well as the vascular recruitment and remodeling by neurons are known to be mediated by soluble factors^3, 5^, it is expected that human endothelial derived soluble factors would also encourage differentiation of the hGPCs to oligodendrocytes. The outcome of interaction between human glial progenitor and human endothelial cells is not known, while subtle inter-species differences are becoming recognized^21^. The potential roles of diffusible factors of human vascular origin in survival, proliferation and differentiation of human glial progenitor cells in context of satem cell niches are therefore a pertinent objective of study specially with the view of the application of the hGPCs in translational therapies. The *in vitro* differentiation fate of GPCs is known to be determined by the composition of culture medium^22^. This study addresses these aspects using human umbilical vein endothelial cells (HUVECs) in the trans-well co-culture system with platelet-derived growth factor receptor-alpha (PDGFRα) sorted bi-potential glial progenitors of fetal human forebrain and thus extends the findings in murine systems to the human glial progenitor cells.

## Results

### HUVECs encourage proliferation of hGPCs under proliferative conditions

PDGFRα sorted hGPCs were maintained under proliferative conditions in co-culture with HUVECs on trans-well in a ratio of 1.2:1 for a period of 4 weeks along with no-co-culture controls in quadruplicate wells with trans-well inserts being changed every 5-7 days. The hGPC spheres originated as adherent spheres by clonal expansion and a sphere visually perceived to be consisting of approximately eight cells was considered as a minimum sphere. Past three weeks in culture the hGPC spheres were found to grow too large to be held to the substrate and they would detach and stay close to the surface of the plate. At this stage the spheres also showed a tendency to gather and fuse together forming mega-spheres. However there was always a significant difference between the number of spheres and the diameter of the largest hGPC sphere between the co-cultured and the no-co-culture controls (Figure 1A). When the number of spheres in individual wells was counted and the counts were analyzed by two way repeated measures ANOVA followed by Bonferroni’s correction of p-values, the co-culture effect was extremely significant (*P* <0.0001) and the increase in sphere numbers was also a function of time (*P* <0.0001) (Figure 1B). In the same experiment, when diameter of the largest sphere was measured, every week for wells with and without HUVEC co-culture, the co-culture effect was extremely significant, with *P* value <0.0001 and along with the time effect with *P* value <0.0001 (Figure 1C). Thus it was concluded that co-culture with the HUVECs encouraged survival or proliferation or both for the PDGFRα positive hGPCs from the fetal forebrain.

**Figure 1.**
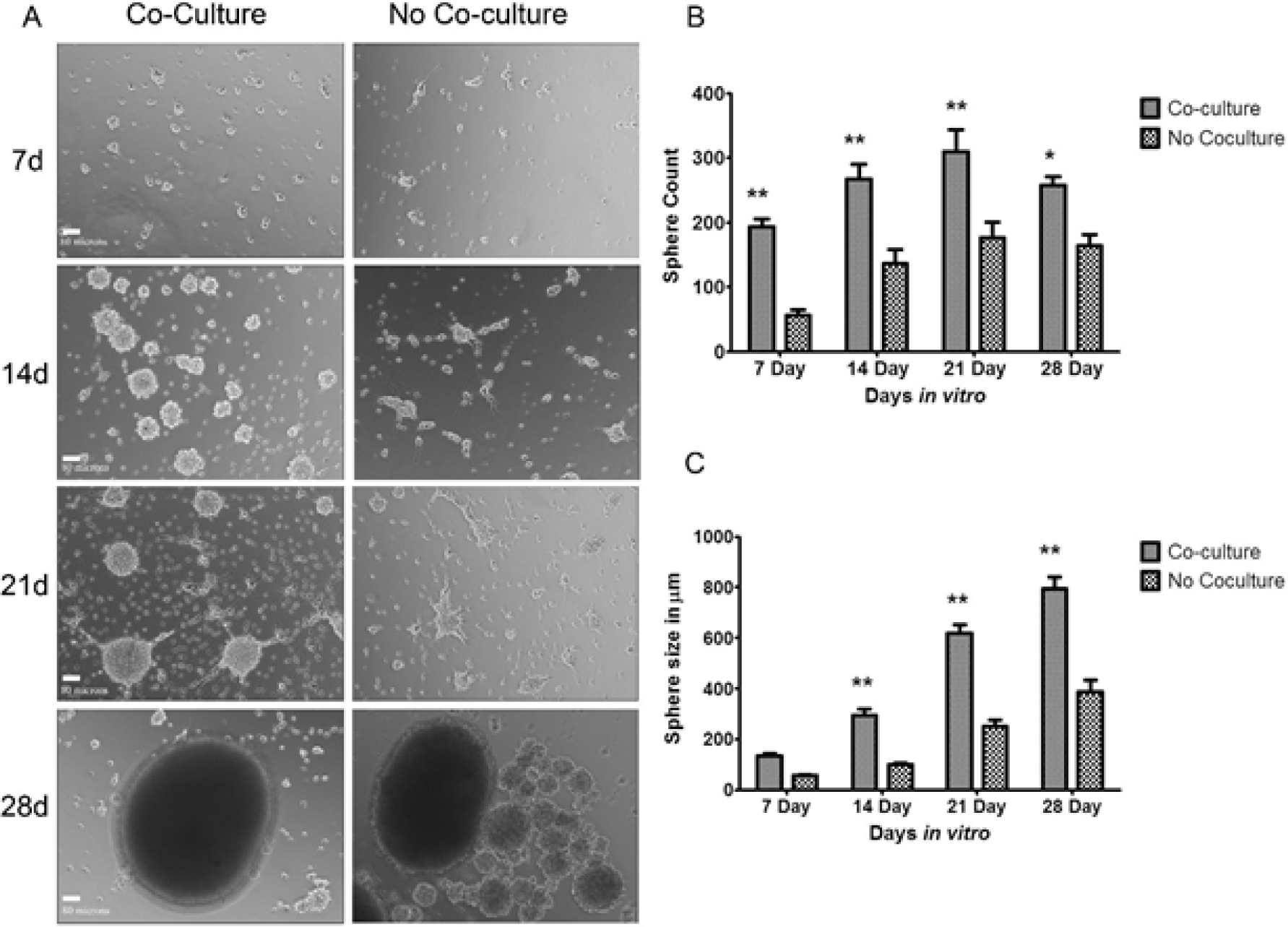
Co-culture with HUVECs stimulates hGPCs sphere formation in a time dependent manner. (A) Representative micrographs of spheres formed by PDGFRα sorted hGPCs from a 22gw specimen in proliferative condition at 7, 14, 21 and 28 days, with or without HUVEC co-culture. Scale bar, 80μm. (B) Average number of spheres formed at 7, 14, 21 and 28 days as mean ± SE from n=3 independent experiments (biological replicates) each with quadruplicate wells for with or without HUVEC co-culture. (C) Average size of the largest sphere hGPC sphere in each well at 7, 14, 21 and 28 days as mean ± SE from n=3 independent experiments (biological replicates) each with quadruplicate wells for with or without HUVEC co-culture. Post hoc p value designated as * for *p* < 0.05 and ** for *p* < 0.01.

### Increased Thymidine Incorporation in hGPCs on co-culture with HUVECs in proliferative conditions

To determine if co-culture with HUVECs encouraged survival and proliferation of the glial progenitors, the hGPC spheres were allowed to grow for up to 4 weeks with or without co-culture and cell numbers in the hGPC spheres were determined by dissociating them with papain/DNase at 7, 14, 21 and 28 days. The effect of HUVEC co-culture on the increase in hGPC cell number was significant with *P* value = 0.0013 with the difference in cell numbers being significant at 21 day and 28 day time points with Bonferroni’s post-test values of *t*(6.208), *p*<0.001 and t(8.991), p<0.001 respectively. Independent of the co-culture, the cell numbers also increased as a function of time with *P* value <0.0001 (Figure 2A). Since increased proliferation should be accompanied by synthesis of new chromatin material, to assess the net proportion of active DNA synthesis in the hGPCs, cells in quadruplicate wells with or without co-culture were pulsed with [^3^H-methyl], thymidine at 4, 12, 18 and 25 days, for 72 hours and thymidine incorporation in the hGPCs was determined. The results showed that the effect of co-culture was significant with *P* value = 0.0052, its effect being significant at only 21 and 28 days with *t* (3.995), *p*<0.01 and *t* (8.819), *p*<0.001 by Bonferroni’s post-tests and not at 7 and 14 days. The effect of time was also extremely significant with *P* value <0.0001 (Figure 2B).

**Figure 2.**
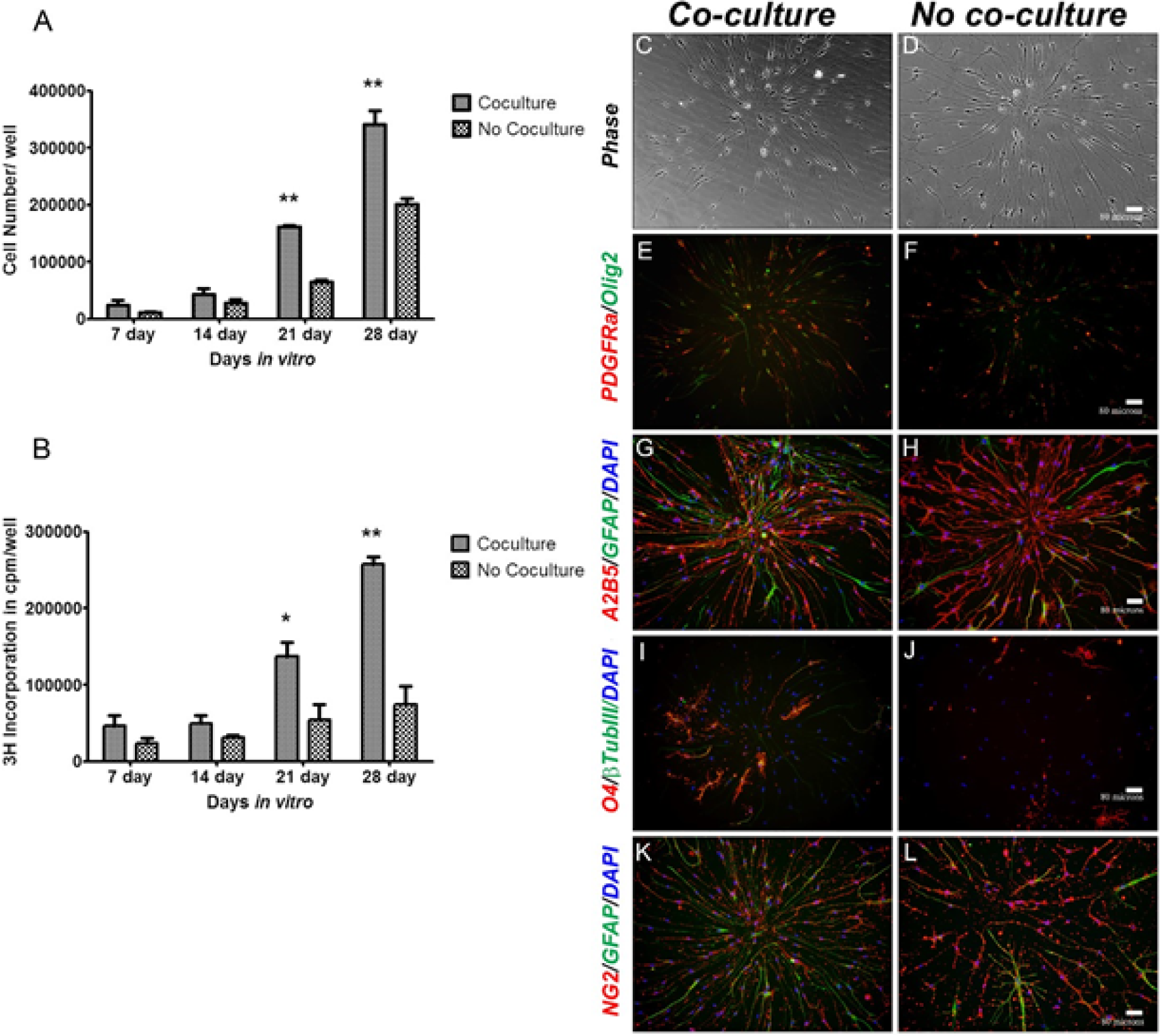
Co-culture with HUVECs induces proliferation of hGPCs but does not alter differentiation outcome of hGPC spheres. (A) Average number of cell count from dissociated hGPC spheres at 7, 14, 21 and 28 days as mean ± SE from n = 3 independent experiments (biological replicates) each with quadruplicate wells for with or without HUVEC co-culture. (B) Average cumulative [3H Methyl] –Thymidine incorporation measured as mean ± SE cpm/well at 7, 14, 21 and 28 days from n=3 parallel independent assays (biological replicates) in replicate quadruplicate wells for with or without HUVEC co-culture. Post hoc p values designated as * for *p*<0.05 and ** for *p*<0.01. (C) Representative images of hGPC spheres from co-culture at 14 days in vitro from a 20.6 gw specimen and allowed to differentiate on poly ornithine – laminin and stained for PDGFRα/Olig2, A2B5/GFAP, O4/beta III tubulin, NG2/GFAP revealed the pattern to be similar for spheres grown with or without co-culture. Nuclear stain is DAPI. Scale bar, 80μm.

### Co-culture in proliferative conditions does not alter differentiation outcome of hGPC spheres

The hGPC spheres formed at the end of each week of culture in proliferative conditions with or without co-culture with HUVECs were re-plated in differentiation conditions and stained for common progenitor and differentiation markers - PDGFRα, A2B5, NG2, Olig2, O4, GFAP and β III tubulin (Figure 2C). The distribution of these markers was not significantly different between the spheres arising from the co-culture and the no co-culture conditions although there were always minor differences in the distribution of markers between sphere to sphere within the same condition. Thus irrespective of co-culture, there were spheres that were remarkably rich in O4 or GFAP compared to the other spheres in the same well. This indicated preservation of asymmetric division of the progenitors under proliferative conditions irrespective of co-culture. Also, regardless of co-culture O4 expression and clonal O4 expansion from the PDGFRα sorted OPCs could be seen only upto first two weeks in these experiments, during which some spheres gave rise to more number of O4 positives than others within the same well. At the end of four weeks *in vitro*, irrespective of co-culture, the PDGFRα sorted hGPC spheres upon re-plating were mostly A2B5 positive (50 to 75%) and GFAP positive (70 to 85%), with very little potential (3 to 6%) to give O4 positive cells.

### Co-culture in differentiating conditions alters differentiation pattern of PDGFRα sorted hGPCs

In order to examine if co-culture with HUVECs could affect the differentiation pattern of progenitors independent of their proliferation, PDGFRα sorted hGPCs were plated directly to differentiating conditions. The cells were fixed at the end of 4 and 7 days and immunocytochemistry was performed for differentiation markers O4 and GFAP as markers for oligodendrocytes and astrocytes respectively (representative micrographs in Figure 3A). The percent of oligodendrocytes as determined by O4 staining decreases significantly upon co-culture with *P* value = 0.0351 in two way ANOVA, with the decrease in the O4 percentage at 7 day being significant with *t* (2.730), *p* < 0.05 while the difference at 4 day being of no significance by Bonferroni’s post-tests (Figure 3B). Time had no significant effect on the O4 survival within this one week. On the other hand, co-culture very significantly caused an induction of astrocytic lineage as measured by the percent of GFAP positive cells, *P* value = 0.0082, with a 26% astrocytic induction at 4 day post-test, *t* (2.894), *p*<0.05 and 31.66% at 7 day with post-test, *t*(3.500), *p*<0.01. Other than the co-culture, the GFAP induction in these cultures was also found to significantly increase with time, with *P* value = 0.0001 (Figure 3C). Thus co-culture with HUVECs encouraged astrocytic differentiation of the PDGFRα sorted hGPCs with a concomitant decrease in differentiation to oligodendrocytic lineage.

**Figure 3.**
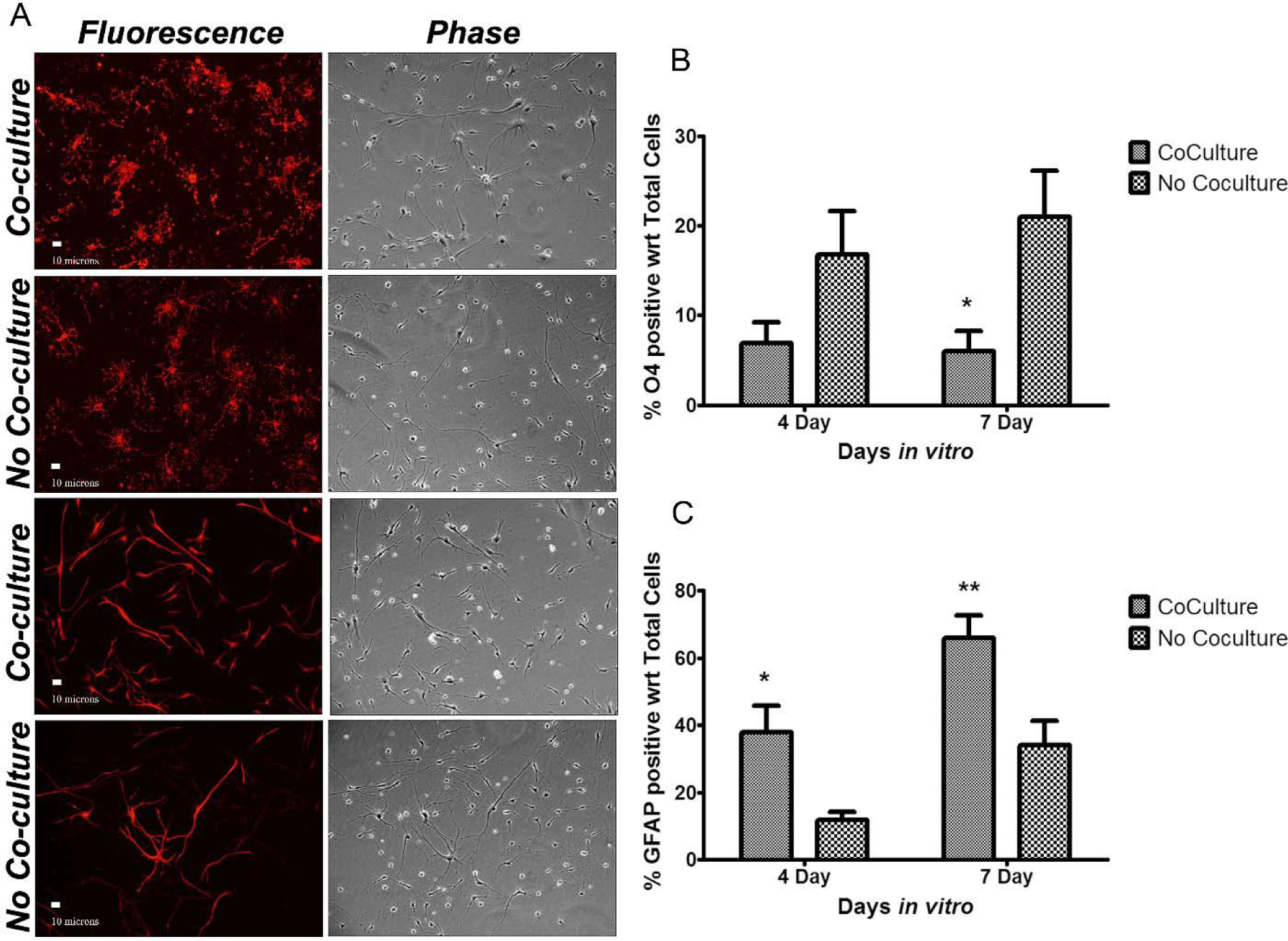
Co-culture with HUVECs induces astrocytic differentiation in the hGPCs. (A) Representative micrographs of hGPCs differentiated on poly-ornithine laminin substrate in presence of 25ng/ml of VEGF in serum free media (20 gw specimen) stained for O4 and GFAP, with or without HUVEC co-culture. Scale bar, 10μm. (B) Percent of O4 positive and (C) Percent of GFAP positive cells out of total cells counted in phase in ten random fields or 200 cells/well as mean ± SE from n=5 independent experiments (biological replicates) each with triplicate wells for with or without HUVEC co-culture. Post hoc p values designated as * for *p*<0.05 and ** for *p*<0.01.

### Transcriptional analysis of HUVECs and hGPCs reveals Bone Morphogenic Proteins, Endothelin1, Gas6 and IL6 as possible soluble mediators of HUVEC influence

In order to dissect the respective contribution of soluble factors from HUVECs and hGPCs to the co-culture environment that may lead to the observed increase in survival, proliferation and astrocytic differentiation of hGPCs in co-culture environment, expression levels of selected relevant genes for soluble factors were assessed by Inventoried Taqman’s human Gene Expression Assay (Inventoried TaqMan® Gene Expression Assays IDs given in Table S1). The HUVEC (n=3) to hGPC (n=4) fold ratio for gene expression with respect to respective endogenous GAPDH in triplicate wells were determined using delta C_T_ method (Table 1, Figure 4A). Of all the 15 genes tested for, the highest difference in mRNA expression relative to GAPDH was found to be for bone morphogenic protein 4 (*BMP4*) which was 134.9 (*p*=0.0026) fold more in the HUVECs than in the hGPCs. Also, bone morphogenic protein 6 (*BMP6*) of transforming growth factor - beta family was found to have a significant 4.211 (*p* = 0.00071) fold more transcripts in HUVECs than hGPCs. Other humoral factor mRNAs that were present in significantly higher abundance in HUVECs were endothelin1 (*EDN1*), a potent vasoconstrictor from endothelial cells with known effects on the central nervous system, with HUVEC to hGPC relative transcript ratio of 41.50, *p* = 0.00101. Still other soluble factors whose transcripts were detected in higher relative abundance in the endothelial cells were growth arrest specific 6 (*GAS6*) which showed 3.753 fold (*p* = 0.02363) and interleukin6 (*IL6*) with 3.726 fold (*p* = 0.00296) higher expression in HUVECs than hGPCs (Figure 4A, Table 1). Also observed were transcripts of humoral factors that were more abundant in hGPCs compared to HUVECs. First among these is Pleiotrophin (*PTN*) which showed a high negative ratio of 0.003612, *p* = 0.01972 followed by fibroblast growth factor 9 (*FGF9*) or glia activating factor with relative ratio of 0.004552 with *p* = 0.02482 for abundance in HUVEC to hGPC. Following them were cytokines, ciliary neurotrophic factor (*CNTF*) and leukemia inducing factor (*LIF*) which were both more abundant in hGPCs than in HUVECs, with a relative ratio of 0.032551, *p* = 0.00710 and 0.048426 with *p* = 0.00235 respectively (Figure 4A, Table 1). Other genes that were assayed and showed expression but did not show differential expression between the HUVECs and the hGPCs were the other members of the TGF beta family, the bone morphogenic proteins 2, 5 and 7 (*BMP2, 5* and *7*) and the fibroblast growth factor -2 (*FGF2*) (Figure 4A, Table 1). Thus HUVECs express a wide variety of soluble factors that are capable of affecting the survival, proliferation and differentiation of the hGPCs. Raw Ct values for these experiments can be seen in Supplemental Spreadsheet 1.

**TABLE 1.** Relative mRNA Expression Fold Ratio by Gene Expression Assay ^a^.

| HGNC<br>Gene<br>Symbol | HGNC<br>Full Name | Entrez<br>Gene<br>ID | HUVEC to hGPC<br>Fold Ratio | Significantly More<br>Abundant in |  | *p*<br>values<br>(n=7) |
| --- | --- | --- | --- | --- | --- | --- |
|  |  |  |  | HUVECs<br>(n=3) | hGPCs<br>(n=4) |  |
| *Bone Morphogenic Proteins* |  |  |  |  |  |  |
| *BMP2* | bone morphogenetic protein 2 | 650 | 0.628235 |  |  | 0.19476 |
| *BMP4* | bone morphogenetic protein 4 | 652 | 134.9001 | Yes |  | 0.00264 |
| *BMP5* | bone morphogenetic protein 5 | 653 | 0.023848 |  |  | 0.10684 |
| *BMP6* | bone morphogenetic protein 6 | 654 | 4.211436 | Yes |  | 0.00071 |
| *BMP7* | bone morphogenetic protein 7 | 655 | 0.128890 |  |  | 0.07468 |
| *Fibroblast Growth Factors* |  |  |  |  |  |  |
| *FGF2* | fibroblast growth factor 2 | 2247 | 1.747718 |  |  | 0.22904 |
| *FGF9* | fibroblast growth factor 9 | 2254 | 0.004552 |  | Yes | 0.02482 |
| *Cytokines* |  |  |  |  |  |  |
| *IL6* | interleukin 6 | 3569 | 3.726636 | Yes |  | 0.00296 |
| *LIF* | leukemia inhibitory factor | 3976 | 0.048426 |  | Yes | 0.00235 |
| *PTN* | Pleiotrophin | 5764 | 0.003612 |  | Yes | 0.01972 |
| *Other Humoral Factors* |  |  |  |  |  |  |
| *GAS6* | growth arrest-specific 6 | 2621 | 3.753718 | Yes |  | 0.02363 |
| *EDN1* | endothelin 1 | 1906 | 41.49811 | Yes |  | 0.00101 |
| *CNTF* | ciliary neurotrophic factor | 1270 | 0.032551 |  | Yes | 0.00710 |
| *Endothelin Receptors* |  |  |  |  |  |  |
| *EDNRA* | endothelin receptor type A | 1909 | 0.045506 |  |  | 0.08647 |
| *EDNRB* | endothelin receptor type B | 1910 | 0.011111 |  | Yes | 0.03155 |

Note: <sup>a</sup> For the expression assay, 5ng of cDNA for each HUVEC (n=3) and hGPC (n=4) sample was loaded in triplicate wells for the Inventoried TaqMan® Gene Expression Assays, the Assay IDs for which are described in Table S1. For each gene, the HUVEC to hGPC expression fold ratio was determined by delta C<sub>T</sub> method after normalizing the expression to endogenous GAPDH. Statistical significance was determined by t test for samples with equal variance. *p*<0.05 was considered significant.

**Figure 4.**
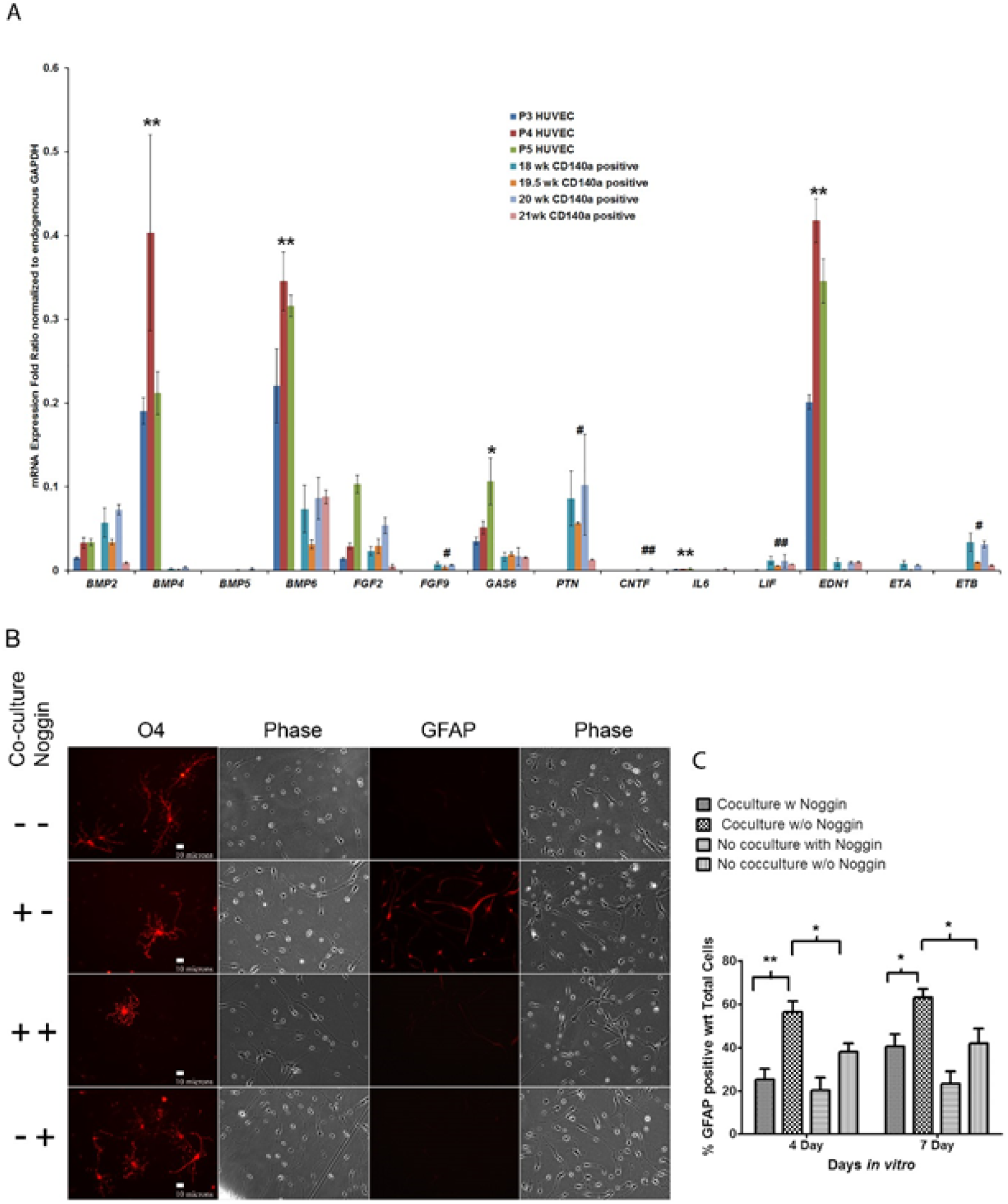
Astrocytic differentiation induced by HUVECs is mediated by soluble factors, particularly the Bone Morphogenic Proteins. (A) Relative mRNA expression fold ratio as mean ± SD for Inventoried TaqMan® Gene Expression Assays IDs given in Table S1 with respect to endogenous GAPDH for triplicate assays (technical replicates) for HUVEC (n=3 biological replicates) and hGPC (n=4 biological replicates). Statistical significance by student’s t-test for equal variance indicated as* for *p*<0.05 and ** for *p*<0.01, when the expression is significantly higher in HUVECs with respect to hGPCs; and as # for *p*<0.05 and ## for *p*<0.01 when the expression is significantly higher in hGPCs with respect to HUVECs as given in Table 1. For raw Ct values please see Supplemental Spreadsheet 1. (B) Representative micrographs for attenuation of HUVEC mediated astrocytic differentiation of PDGFRα sorted hGPC by 100ng/ml Noggin (19.3 gw specimen). Scale bar, 10μm. (C) Percent of GFAP positive cells after treatment with 100ng/ml Noggin out of total cells counted in phase in ten random fields or 200 cells/well as mean ± SE from n=4 independent experiments (biological replicates) each with triplicate wells for with or without HUVEC co-culture reveals inhibition of astrocytic differentiation of hGPCs. Post hoc p values designated as * for *p*<0.05 and ** for *p*<0.01.

### Noggin can inhibit HUVEC induced Astrogenesis

Based on the relative abundance ratios of the transcripts seen in gene expression analysis, bone morphogenic protein 4 was expected to be the major humoral factor from HUVECs in co-culture media that could affect the differentiation fate of hGPCs in co-culture. Thus it was recognized that if the observed astrocytic differentiation of the PDGFRα sorted hGPCs in co-culture with HUVECs is determined by the BMPs, the same should be attenuated by the general inhibitor of BMP signaling - Noggin. Therefore, PDGFRα sorted hGPCs were directly plated for differentiation with a pre-treatment by 100ng/ml of Noggin for 4h to block all endogenous BMP signaling followed by start of co-culture with HUVECs. The Noggin treatment was repeated every 24 hours till 4 and 7 days followed by immunocytochemistry for O4 and GFAP respectively for assessment of commitment to oligodendrocyte or astrocyte lineage respectively (Figure 4B). Statistical analysis of the counts by one way ANOVA revealed that Noggin could effectively reduce HUVEC induced astrogenesis at both 4 and 7 day time points with *P* = 0.0012 with an effective lowering of the percent of GFAP positive cells from 56.27 ± 5.085 (mean ± S.E.) for co-culture without noggin to 25.21 ± 4.814 for co-culture along with noggin, with *p* < 0.01. On the other hand only 37.83 ± 4.042 percent of hGPCs showed GFAP expression in absence of co-culture under identical conditions, *p* < 0.05. At day 7, the daily noggin treatment continued to attenuate HUVEC mediated astrogenesis with similar statistical pattern (Figure 4C). Since HUVEC mediated astrogenesis could be completely attenuated by noggin, the effect was concluded to be primarily BMP mediated.

### Network Analysis also reveals BMP4 as the key differentiation factor

Since transcripts of soluble factors other than BMP4 over-expressed by HUVECs relative to hGPCs can also make significant contribution towards the observed astrocytic differentiation, molecular networks based on genetic and protein interactions with gene lists specific to the co-culture and no co-culture environments (Input Gene-lists for Differential Networks given in Table S2) were used to understand their relative contributions. Networks built on PCViz2 application of Pathway Commons using the respective gene lists were analyzed on Cytoscape 3.4.0 to generate subnetworks for the no co-culture and co-culture conditions which were represented with statistical weightage of molecular interactions at the nodes as well as the edges (Figure 5A & B). The difference between the two networks generated for co – culture and no co-culture environments (Figure 5C) revealed BMP4 to be the node with most number of connections, followed by EDN1, BMP6 and IL6 while GAS6 was the least connected node in this context. This closely follows the earlier observation of inhibition HUVEC induced of astrocytic differentiation of PDGFRα sorted hGPCs by Noggin and further validated the predominance of BMP4 in this context. Complete list of nodes and edges for all three networks can be accessed from Supplemental Spreadsheet 2.

**Figure 5.**
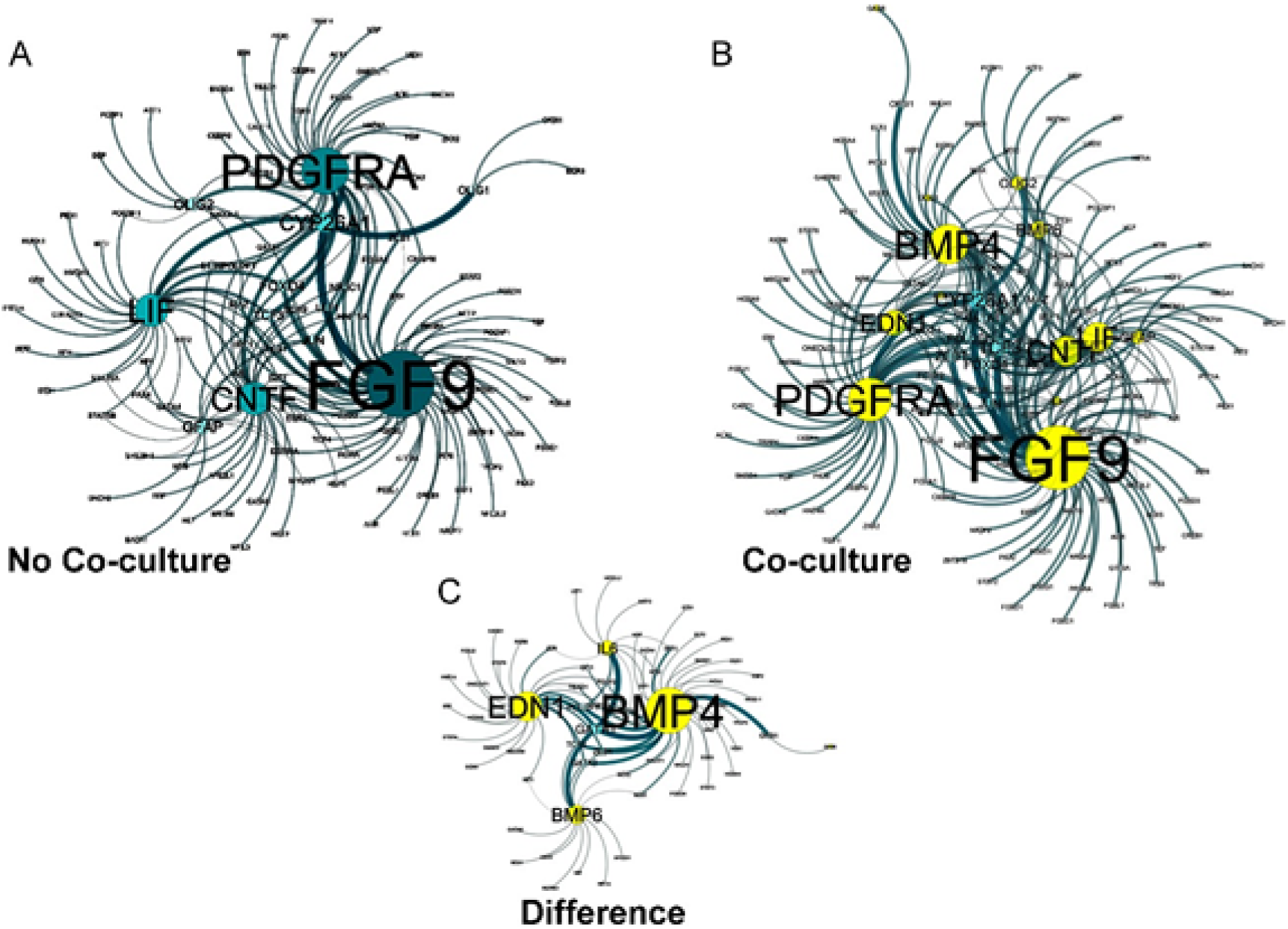
Molecular Network Analysis confirms BMP4 to be the key soluble factor behind astrocytic differentiation. Networks were generated for no co-culture and with co-culture condition using Input Gene-lists for Differential Networks given in Table – S3 on the PCViz2 application of Pathway Commons which were visualized on Cytoscape 3.4.0 for identification of major subnetworks and generation of difference network (A) Network generated for the No Co-culture condition. (B) Network generated for the co-culture condition (C) Difference Network of the Co-culture and the No co-culture condition. In all networks circles are larger for nodes with higher connectivity and broader edges indicate stronger connections. In network (B) and thus also in (C) nodes indicated by yellow circles are those provided in the input gene list. For complete list of nodes and edges for all three networks, see Supplemental Spreadsheet 2.

## Discussion

The study focuses on the outcome of human endothelial co-culture on PDGFRα sorted hGPCs particularly from the perspective of the effect of endothelial derived humoral factors by using a trans-well co-culture model since it is expected to have significant implication towards the application of these hGPCs in translational therapies. The study confirms that human umbilical vein endothelial cells (HUVECs) can affect the biology of PDGFRα sorted fetal human glial progenitor cells (hGPCs) by encouraging their survival, proliferation and differentiation towards astrocytes and bone morphogenic proteins, possibly BMP4, are the primary mediator of this specification towards astrocytic lineage. BMP6, endothelin1, IL6 and Gas6 could be the other possible mediators of these events. The results are in agreement with previous reports implicating BMPs as inhibitors of oligodendrocyte development and promoters of astrocyte development in cultures of embryonic precursor cells from murine cerebral cortex, with the BMP inhibitor noggin having the opposite effect^19,20,23^. The study confirms those findings to be true for the PDGFRα sorted fetal hGPCs in co-cultures with endothelial cells of human origin. Induction of astrocytic differentiation by endothelial cells has been long known^16,17,24^, however this is the first report of demonstration of the same for the PDGFRα sorted fetal hGPCs when both the endothelial and the glial progenitor cells in co-culture are of human origin. BMP signaling has been identified as one of the key regulatory pathway for self-maintenance and differentiation for A2B5 sorted glial progenitor which also overexpress BMP4 inhibitors, neuralin and BAMBI (BMP and activin membrane-bound inhibitor) suggesting tonic defense against BMP signaling^25^. The present study indicates that the inherent tonic defense is insufficient in presence of the excess of BMP4 presented by the endothelial cells during co-culture which successfully overrides it to induce astrocytic differentiation of the PDGFRα sorted hGPCs. Leukemia inhibitory factor (*LIF*) and ciliary neurotrophic factor (*CNTF*) whose transcripts were also determined to be relatively more abundant in hGPCs are also known to promote astrocytic differentiation of GPCs to progenitor astrocytes and mature astrocytes^26,27^.

The gestational age of the fetus from which the hGPCs are derived can be ruled out from being a confounding factor behind these observations because all progenitors have been derived from 18-23gw. It is known that A2B5- and PDGFRα-selected progenitors derived from 15-23gw exhibit an age-dependent increase in early to mid-stage lineage markers, including sulfatides (O4 antibody) and transcription factor Olig2^28^. In this study, HUVECs have been used as the endothelial partners in co-culture as they are readily available, most commonly used source of primary endothelial cells and widely described. Comparative expression profiles of HUVECs and human cerebral endothelial cells (HCEC) are available^29^ where IL6 has been reported to be exclusively released by HCEC and not HUVECs. However in this study IL6 transcripts were found to be more abundant in HUVECs than in hGPCs, making IL6 one of the diffusible factors that could contribute to the observed effects of co-culture. Although it is known that BMP4 accelerates the commitment of human embryonic stem cells (ESCs) to the endothelial lineage^30^, influence of hGPC’s on HUVEC proliferation and maintenance was outside the scope of this study. However it can be noted that, other than the abundance of BMP transcripts in the HUVECs themselves, relative abundance of the pleiotropin (*PTN*) transcripts in the hGPCs, a known factor in neovascularization^31^, opens possibilities of existence of a positive feedback loop between these two cell types. In a relatively fresh approach, a comparative of transcription profiles of a selection of diffusible factors from the two cell types – HUVECs and hGPCs - relevant to the context of niche environment has been used to establish their comparative significance.

PDGF signaling activation has been widely implicated in brain tumors^32, 33^ and PDGFRα positive progenitor cells have been associated with the early changes associated with tumor initiation^34^. PDGFRα expressing stromal cells derived from oligodendrocytes progenitor cells (OPC) have been discovered at the invasive front of high-grade gliomas, exhibiting an unique perivascular distribution^35^ indicating that the interaction of these cells with the endothelial partners in the tumor niche could be of significance towards tumor origin or expansion. Though BMP4 is generally understood to be involved in differentiation pathways by a reduction in proliferation^36^, there are also reports of its being involved in cell proliferation and maintenance of the pluripotent state of mouse ESCs^37^. During the present study, when the effect of noggin on HUVEC mediated proliferation of hGPCs was studied with a pre-treatment of hGPCs with 100ng/ml of noggin for 4h followed by daily treatment with noggin every 24h for upto 14 days in wells with or without co-culture as described in Supplemental Experimental Procedures. hGPC sphere count and cumulative thymidine incorporation assay revealed that noggin could effectively bring down the proliferation of hGPCs in co-culture with HUVECs compared to no co-culture controls. Thus, BMPs do play a role in proliferation of the hGPCs in this context, aside the PDGF-AA and FGF2 which are already present in the proliferative media.

Although endothelin1 has been demonstrated to signal oligo progenitor migration and differentiation in postnatal brain of mice^38^, endothelin axis is also known to be involved in many human malignancies and endothelin receptor antagonists have anti-tumorigenic potential^39^. Therefore, when endothelin1, was detected as the second most abundant transcript of soluble factor from the HUVECs, level of expression of its receptors - type A (*EDNRA*) and type B (*EDNRB*) were also assayed in both the cell types to envisage its possible role in the context. In the gene expression assays, *EDNRA* did not show any significant difference in expression in the hGPCs with respect to the HUVECs while *EDNRB* was significantly more abundant in PDGRFα sorted hGPCs compared to HUVECs (Figure 4A, Table 1).

Endogenous expression of *PTN* by the hGPCs as observed in this study has also been reported previously for adult WMPCs^25^ and it increases the tumorigenic potential of hGPCs as PTN is associated with tumor angiogenesis^31,40^. Additionally, GAS6 a survival factor for hGPCs was found to be one of the more abundant transcripts expressed by the HUVECs. AXL the cognate receptor for GAS6, which are known candidate genes for glioma therapy^41,42^, was also detected in the hGPCs by immunocytochemistry particularly more when the hGPCs are astrocytic or progenitor like than oligodendrocytic in morphology (Figure S1). This indicates, any GAS6 mediated survival benefit is more likely to be effective when the cells in progenitor or astrocytic form. IL6 the other soluble factor transcript abundant in the HUVECs is also a known tumor promoting factor, targeting which is known to suppress glioma stem cell survival and tumor growth^43^.

Thus exposure to almost all soluble factors relatively more prevalent in the endothelial cells increases tumorigenic potential of the PDGFRα sorted fetal hGPCs than their oligogenic potential in absence of extracellular matrix (ECM) anchorage. This is in complete contradiction to the premise by which soluble factors from endothelial cells were expected to encourage oligodendrocytic differentiation in the same lines as they encourage neuronal differentiation mediated by BDNF^3,5^. However anatomy of stem cell niches encourages stem cells to be in close contact with laminin containing ECM surrounding vascular endothelial cells. One of the functional roles for α6β1 integrin, the laminin receptor on stem cells is inhibiting the stem cell proliferation through ECM interaction within the vascular niche^1^. Also adhesion to vasculature is the key to directed migration of the OPCs before maturation^18^.

The understanding derived from this study of PDGFRα sorted hGPCs in co-culture with HUVECs on trans wells replicates an *in vivo* situation where hGPCs are in close enough vicinity to interact with endothelial derived soluble factors but do not get exposed to the anti-proliferative signals from ECM. This is likely to encourage undesired proliferation as observed for the PDGFRα sorted hGPCs in this study. Thus although successful re-myelinations are being observed with the PDGFRα sorted hGPCs in the hypomyelinated shriverer mouse brain^11^, there remains still the need to define a stricter unipotential oligoprogenitor lineage that would mature to only myelinating cells on transplantation. One emerging candidate in this regard are the PDGFRα positive cells that also co express the pro-oligodendroglial tetraspanin CD9^11,44^. The study suggests, loss of contact of the transplanted PDGFRα sorted hGPCs progenitors with vasculature ECM might lead to their uncontrolled expansion induced by the humoral factors from the niche endothelial cells. Therefore, caution needs to be maintained to avoid possibilities of tumorigenesis while recommending transplantation therapies with these cells for ensuring safety of the translational procedures and unipotent progenitors for oligodendrocytes needs to be further investigated.

## Materials and Methods

### Cells

#### Platelet Derived Growth Factor Receptor alpha (PDGFRα) positive human Glial Progenitor Cells (hGPCs)

All experimental protocols were approved by the University of Rochester – Strong Memorial Hospital Research Subjects Review Board. All methods were carried out in accordance with the approval and relevant guidelines and regulations. Informed consent was obtained for tissue usage from aborted fetuses used in this study between 18 to 2weeks in gestation. All of the 15 fetal brain specimens used in this study were from aborted fetuses between 18-23 gestational weeks and were obtained after cases of abortion, with informed consent to tissue use under protocols approved by the University of Rochester – Strong Memorial Hospital Research Subjects Review Board. The cortical specimens were collected and dissociated within less than 2h post mortem using papain and DNase as per established protocols and the cells were plated at 2 million/ml in serum free media (SFM) containing DMEM/F12/N1 (Complete SFM composition for hGPCs in Table S3) with 20 ng/ml of fibroblast growth factor 2 (FGF2) for overnight. The dissociated cells were recovered the following day, counted and sorted for Platelet Derived Growth Factor Receptor alpha (PDGFRα) positive hGPCs, identified using the primary antibody against an external epitope of PDGFRα, CD140a, (Cat#556001, BD Pharmingen) and rat secondary anti-mouse IgG2a+b conjugated Microbeads (Order No.130-047-201, Miltenyi Biotech) by magnetic cell sorting^11^. After sorting, cells were either frozen for isolation of total mRNA or seeded in SFM as defined for proliferative or differentiating conditions in co-culture with HUVECs.

#### Human Umbilical Vein Endothelial Cells (HUVECs)

The human Umbilical Vein Endothelial Cells were purchased from the Department of Microbiology and Immunology, University of Rochester Medical Center in their early (second) passage in M-200, Low Serum Growth Supplement and 5% FBS (Cascade Biologicals) and were maintained on 0.2% gelatin coated tissue culture flasks in DMEM/F12, 10% PD-FBS, 10ng/ml of FGF2 and 25ng/ml of VEGF. The cells were passaged every 5-7 days with 0.05% Trypsin/ EDTA with 1:4 splitting and used from passage 3-5 for total mRNA isolation as well as co-culture with hGPCs.

### Co-culture

For hGPC - HUVEC co-culture, approximately 1:1 ratio (exactly 1.2:1 at plating) was maintained between the two cell types. PDGFRα positive glial progenitors (hGPCs), 17500 in number, were seeded in each well of the 24 well plate directly post sort on defined substrate and SFM for either proliferative or differentiating condition and 15000 HUVECs were seeded on 0.2% gelatin coated trans-well inserts with 0.4μm pore size.

*Proliferative condition* was defined by untreated 24 well suspension plate as substrate hGPCs in SFM with 20 ng/ml of FGF2, 20ng/ml of PDGF-AA for co-culture. The co-culture was carried out for 28 days, with a change in the trans-well insert every 5-7 days up to 28days for the remaining plates. Plates were withdrawn for assessment of hGPC sphere formation each week using protocols detailed in Supplemental Experimental Procedures.

*Differentiating condition* was defined by the presence of poly-ornithine-laminin coat on the 24 well adherent plate as substrate for hGPCs in SFM with 25ng/ml VEGF to support the HUVECs with alternate day change of media. The culture was terminated by fixing the cells as required for immunocytochemistry at 4 and 7 days.

*Inhibition of differentiation.* For inhibition of differentiation with Noggin, the hGPCs were pretreated with 100ng/ml recombinant human Noggin (R & D Systems) 4h prior to start of co-culture to inhibit all endogenous BMP signaling and re-treated every 24h thereafter. At the end of 4 and 7 days, the cells were fixed and immunocytochemistry was performed for O4 and GFAP.

### Immunocytochemistry

Immunocytochemistry was done for the hGPCs at the end of 4 or 7 days of co-culture under differentiating condition or at the end of 2h or 4d after re-plating of the spheres for differentiation, using antibody specific protocols for primary antibodies against PDGFRα, A2B5, NG2, Olig2, GFAP, β–III tubulin, O4, GFAP and Axl as detailed in the Supplemental Experimental Procedures. The differentiation experiment was performed with five times and the inhibition of differentiation performed four times (biological replicates each with triplicate wells for with or without HUVEC co-culture) and counts presented as mean ± SE.

### Thymidine Incorporation Assay

Under proliferative condition, hGPC cultures were pulsed with 1µCi of (Methyl-^3^H) – Thymidine per ml of media in quadruplicate wells with and without co-culture and incubated for 72 h without any further change of media. At the end of incubation the cells were washed with Hanks solution, DNA precipitated with ice-cold 5% (wt/vol) trichloroacetic acid for 20 min at 4°C, and solubilized in 0.55 ml of 0.3 M sodium hydroxide – 0.1% sodium docecyl sulfate for 1– 2 h at 37°C. To 5 ml of Ecoscint A scintillation cocktail, 0.5 ml of this cell extract was added and radioactivity measured in a Beckman Coulter LS 6500 Liquid Scintillation Counter for 1 minute. Experiment was repeated with three biological replicates, in quadruplicate wells each for co-culture and no co-culture and data expressed as mean ± SE cpm /well.

### RNA Preparation and Gene Expression Assay

Total RNA was extracted from thawed PDGFRα positive hGPCs (n=4 biological replicates) and HUVECs (n=3 biological replicates) that had been snap frozen earlier directly post-sort or post-passage respectively, using RNeasy Mini kit (Qiagen) for Animal Cells with “on column” DNase digestion. Total RNA was reverse transcribed using the Reverse Transcription reagents (Applied Biosystems), random hexamers and MultiScribe™ Reverse Transcriptase (RT). Inventoried TaqMan® Gene Expression Assay with 900nM forward and reverse primers and 250nM FAM-labeled MGB probes were performed for each of the 15 relevant genes for soluble factors (Inventoried TaqMan® Gene Expression Assays IDs in Table S2) with 5ng of the cDNA in triplicate wells using TaqMan® Universal PCR Master Mix on a thermal cycle of - 2 min at 50 °C, 10 min at 95 °C, followed by 40 cycles of 15 sec at 95 °C and 2 min at 60 °C on ABI Prism 7000. Human GAPDH (FAM / MGB Probe, Non-Primer Limited) was used as endogenous control and No RT control was performed for each RNA sample for every assay. Gene expression was determined for HUVECs and hGPCs relative to their respective endogenous GAPDH by delta C_T_ method followed by relative abundance ratio of HUVEC to hGPC for the respective gene with significance determined by t-statistics. For raw Ct values please see Supplemental Spreadsheet 1.

### Network Analysis

Differential environment based molecular networks were generated using gene lists relevant to co-culture and no co-culture environment (Input Gene-lists for Differential Networks given in Table S3) consisting of the phenotypic marker genes used in sorting and immunocytochemistry in this study and the soluble factor genes assessed here in the gene expression assay. While both the gene lists contained the same list of phenotypic markers as well as the soluble factors relatively over expressed by the hGPCs, the gene list for co-culture environment in addition also contained the humoral factors found to be over-expressed by HUVECs relative to hGPCs in gene expression assay. Differential networks were generated using these two gene lists on the PCViz2 application of Pathway Commons^45^. Network BioPAX files^46^ were downloaded and opened on Cytoscape 3.4.0^47^ for identification of major subnetworks, difference networks between the co-culture and no co-culture conditions. Statistical visualization of molecular interactions was done to detect the statistically most prevalent soluble factor in the co – culture and the no co-culture environments based on the number of connectivity. For complete list of nodes and edges for all three networks please see Supplemental Spreadsheet 2.

### Statistical Analysis

Statistical Analysis was performed on Prism (v5.02), GraphPad Software, CA. Two way Repeated Measures ANOVA, followed by Bonferroni’s correction of p values was used for differences in sphere count, sphere size, thymidine incorporation, cell numbers post sphere dissociation and the effect of co-culture on O4 and GFAP expression by hGPCs. One way ANOVA followed by Neuman-Keul’s multiple comparison post-test was used for evaluating the effect of Noggin to attenuate astrocytic differentiation of hGPCs in co-culture. Student’s t-test for samples with equal variance was used for determining significance in gene expression assays. *P* < 0.05 was considered statistically significant and data are presented as mean ± SE except for gene expression assays where it is reported as mean ± SD.

## Supporting information

Supplemental Experimental Procedures

## Data Availability

The datasets generated during and/or analyzed during the current study are available from the corresponding author on reasonable request.

## Acknowledgements

The author thanks the University of Rochester for the postdoctoral fellowship from Neurology Fellowship Program for investigative training in Clinical Neuroscience in the area of Cell and Gene Therapy at the Centre of Translational Neuromedicine. NINDS Grants P01NS050315 & R01NS039559 and NRSA Grant F31NS070441 to Goldman Lab supported the work.

## Author Contributions Statement

A.D. was responsible for designing the experiments, executing them and collecting the data.

A.D. also analyzed the data, wrote the first to the final draft of the paper.

