## Supplemental Experimental Procedures for "Endothelial Cells stimulate proliferation of CD140a sorted human Glial Progenitor Cells and their specification towards astrocytic lineage"

**Asmita Dasgupta^1, 2,*^**

**1. Department of Biochemistry and Molecular Biology, Pondicherry University, Pondicherry - 605014, India.**

**2. Formerly, Postdoctoral Research Associate, Centre of Translational Neuromedicine, Division of Neurology, University of Rochester, Rochester, NY 14642, USA.**

**Inventory of Supplemental Information**

**SUPPLEMENTAL DATA**

**Figure S1.** **Expression levels of Axl receptors for GAS6 in hGPCs by Immunocytochemistry.**

**Table S1. Inventoried TaqMan® Gene Expression Assays IDs**

**Table S2. Input Gene-lists for Differential Networks**

**Table S3. Complete Serum Free Media (SFM) Composition for the hGPCs**

**SUPPLEMENTAL EXPERIMENTAL PROCEDURES**

**Assessment of hGPC Sphere formation**

**Immunocytochemistry**

**Sphere inhibition assays**

**SUPPLEMENTAL SPREADSHEETS**

**Supplemental Spreadsheet 1 – Inventoried TaqMan® Gene Expression Assays Results**

**Supplemental Spreadsheet 2 – Network Nodes and Edges**

**SUPPLEMENTAL DATA**

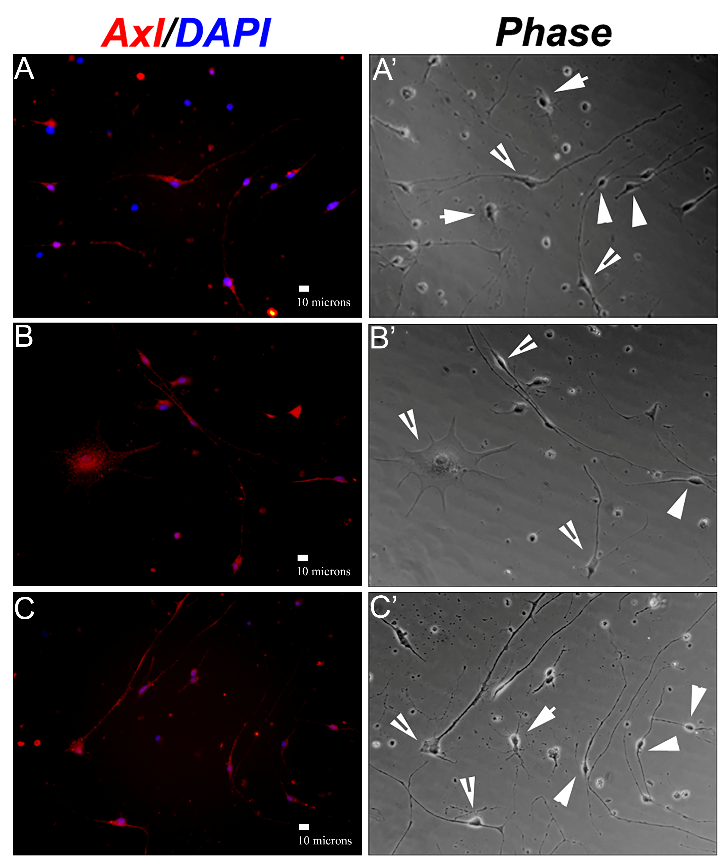

**Figure S1.** **Expression levels of Axl receptors for GAS6 in hGPCs by Immunocytochemistry.** Gas6 was one of the soluble factors relatively overexpressed by the HUVECs than hGPCs and hence the expression of one of its cognate receptor Axl, a receptor tyrosine kinase, was assessed by immunocytochemistry in hGPCs after four days in vitro under differentiation conditions. Primary rabbit anti - Axl - C-terminal polyclonal (at 1:200, ab72069, Abcam) was followed by goat anti rabbit IgG Alexa 568 (1:400) on the hGPCs. It was observed that the hGPCs express Axl in a very differential manner among its heterogeneous phenotypes. The cells expressing most Axl were more astrocytic in morphology followed by moderate staining on the cells with the bipolar progenitor phenotype while the cells with characteristic oligodendrocytic morphology showed least expression for the receptor Axl. Thus in the event of Gas6 acting as a survival and growth factor for the hGPCs the cells most likely to derive advantage from such receptor tyrosine kinase activation would be the cells of the astrocytic lineage or the progenitors themselves while the oligodendrocytes would be least likely to derive much advantage. The representative images (A) – (C) are three random fields in fluorescence and (A’) – (C’) the corresponding images in phase from the same well of a 20 wk sample showing the Axl receptor to be most expressed in cells with astrocytic morphology (indicated by open arrowheads on the phase image) followed by progenitors (indicated by closed arrowheads) while being least expressed by oligodendrocytes (indicated by arrowheads on the matched phase image).

**Table S1. Inventoried TaqMan® Gene Expression Assays IDs ^b^**

| HGNC  Gene Symbol | NCBI Reference  Sequence | Taqman Assay ID | Amplicon Length (bp) |
| --- | --- | --- | --- |
| BMP2 | NM_001200.2 | Hs01055564_m1 | 84 |
| BMP4 | NM_001202.3 | Hs00370078_m1 | 58 |
| BMP5 | NM_021073.2 | Hs00234930_m1 | 67 |
| BMP6 | NM_001718.4 | Hs01099594_m1 | 108 |
| BMP7 | NM_001719.2 | Hs00233476_m1 | 73 |
| FGF2 | NM_002006.4 | Hs00266645_m1 | 82 |
| FGF9 | NM_002010.2 | Hs00181829_m1 | 83 |
| Gas6 | NM_000820.1 | Hs01090305_m1 | 60 |
| PTN | NM_002825.5 | Hs00383235_m1 | 76 |
| CNTF | NM_000614.3 | Hs00173456_m1 | 73 |
| IL6 | NM_000600.2 | Hs00174131_m1 | 95 |
| LIF | NM_002309.3 | Hs01055668_m1 | 77 |
| EDN1 | NM_001955.3 | Hs00174961_m1 | 62 |
| EDNRA | NM_001957.2 | Hs00609865_m1 | 98 |
| EDNRB | NM_000115.2 | Hs00240747_m1 | 77 |
| GAPDH | NM_002046.3 | Hs99999905_m1 | 122 |
| ^b^ Total RNA from the HUVECs post passage and the hGPCs post sort were reverse transcribed and expression for the selected genes were assayed using Inventoried Taqman’s Gene Expression Assay, the Assay IDs for which are presented above. HGNC= HUGO Gene Nomenclature Consortium. | | | |

**Table S2. Input Gene-lists for Differential Networks**

| No Co-culture | With Co-culture |
| --- | --- |
| Phenotypic Markers | |
| *PDGFRA* | *PDGFRA* |
| *OLIG1* | *OLIG1* |
| *OLIG2* | *OLIG2* |
| *GFAP* | *GFAP* |
| Soluble factors | |
| *FGF9* | *FGF9* |
| *PTN* | *PTN* |
| *CNTF* | *CNTF* |
| *LIF* | *LIF* |
|  | *BMP4* |
|  | *BMP6* |
|  | *EDN1* |
|  | *IL6* |
|  | *GAS6* |

**Table S3. Complete Serum Free Media (SFM) Composition for the hGPCs**

| **SFM Component** | **Final Concentration** | **Vendor** | **Catalog #** |
| --- | --- | --- | --- |
| DMEM/F12 | 1X | Invitrogen | 11330-032 |
| HEPES | 17.9mM | Invitrogen | 15630-080 |
| L-Glutamine | 6.35mM | Invitrogen | 25030-081 |
| NEAA | 0.15-0.34mM | Invitrogen | 11140-050 |
| D-Glucose | 46.4mM | Sigma | G8769-100ml |
| Sodium pyruvate | 1.5mM | Invitrogen | 11360-070 |
| Pen/strep | 40U/ml | Invitrogen | 15140-122 |
| BSA | 0.01% | Invitrogen | 15260-037 |
| N1  Insulin  Transferrin  Putrescine | 1X  5µg/ml  5µg/ml  0.0573 µg/ml | Sigma | N6530-5ml |
| Selenite | 0.065 µg/ml | Sigma | S9133-1mg |
| Progesterone | 0.0573 µg/ml | Sigma | P6149-1mg |

**SUPPLEMENTAL EXPERIMENTAL PROCEDURES**

**Assessment of hGPC Sphere formation**

The hGPC sphere formation at 7, 14, 21 and 28 days in quadruplicate wells with or without co-culture for all three biological replicates was monitored by counting and measuring the size of the largest sphere on an Olympus IX70 at 10X (NA 0.3) using an eyepiece grid with hemocytometer markings as reference. At the end of each week the spheres after counting and measuring were dissociated with papain/DNase and number of cells/well was determined for co-culture and non-co-culture conditions. Some hGPC spheres were also re-plated on poly-ornithine-laminin coated substrate with serum free media and processed for immunocytochemistry at the end of 2h or 4 day for following the differentiation pattern of the spheres. On replicate set of plates, cumulative ^3^H-Thymidine Incorporation Assay was performed at 7, 14, 21 and 28 days under the same conditions. All experiments were performed thrice (biological replicates) with quadruplicate wells for both conditions.

**Immunocytochemistry**

*Staining:* For the early oligidendrocyte marker O4, the cells were fixed with 1% PFA, live staining was done for PDGFRα, A2B5 and NG2; while the cells were fixed with 4% PFA for Olig2, GFAP, β–III tubulin and Axl. Primary antibodies were - mouse anti human CD140a monoclonal, IgG2a (anti-PDGFRα) (at 1:100, 556001, BD Pharmingen), mouse anti-A2B5 (neuron cell surface antigen) monoclonal IgM (at 1:500, MAB312, Chemicon), mouse anti-oligodendrocyte marker O4 monoclonal IgM (at 1:250, MAB345, Chemicon), rabbit anti-NG2 chondroitin sulfate proteoglycan polyclonal (at 1:200, AB5320, Chemicon), rabbit anti-Olig2 polyclonal (at 1:200, ab33427, Abcam), mouse anti GFAP monoclonal IgG1 (at 1:1000, SMI 21, Sternberger Monoclonals), mouse anti-Neuronal Class III β-Tubulin monoclonal IgG2a (at 1:1000, MMS-435P, Covance), rabbit anti - Axl - C-terminal polyclonal (at 1:200, ab72069, Abcam). The staining was visualized using species and isotype specific Alexa Fluor (Alexa 488, Alexa 568 and Alexa 594) tagged goat secondary antibodies at 1:400.

*Microscopy:* All visualization, counting and imaging was done using Olympus IX70 Inverted Microscopes from Olympus Corporation. For the differentiation and inhibition of differentiation experiments, number of cells positive for O4 or GFAP were counted on ten random microscopic fields of approximately uniform density to a total of 200 cells for each well and the percent of O4 or GFAP positive cells were expressed out of the total number of cells counted on phase in that well. All counting was done on 20X objective (NA 0.7) across 10 random fields of approximately uniform density for each well. Sphere diameters were measured using relative measurements of standard hemocytometer grids aligned to eyepiece grids on the Olympus IX70. The sphere images were taken at a magnification of 10X (NA 0.3) on the objective using a Microfire® real-time progressive scan true color microscope camera while the immunofluorescence images were taken using 10X (NA 0.3, for the sphere immunocytochemistry) , 20X and 40X (NA 0.7 and 1.15 respectively, for the rest of the images) with an Olympus DP70 Digital Microscope Camera. Exposure values were matched between the experimental and control during imaging for every single antigen. For multiple staining on the same field the color images were merged directly using the DP70 camera software to maintain their integrity. Adobe Photoshop CC was subsequently used for all image data processing to their final publication form.
